# SMECT: a framework for benchmarking post-GWAS methods for spatial mapping of cells associated with human complex traits

**DOI:** 10.64898/2026.02.14.705903

**Authors:** Meng Liu, Chao Xue, Ying Luo, Wenjie Peng, Lihang Ye, Liubin Zhang, Wu Wei, Miaoxin Li

## Abstract

Spatially resolving the cellular basis of complex human traits is essential for elucidating disease mechanisms, yet the comparative performance of computational methods for this task has not been systematically evaluated. Here, we present SMECT (Spatial Mapping Evaluation of Complex Traits), the first comprehensive framework for systematically evaluating methods that integrate genetic data with spatial transcriptomics. SMECT combines a biologically realistic simulation engine, a curated resource of 21 diverse real-world datasets, and a multi-faceted assessment toolkit. Using this framework, we benchmarked three state-of-the-art methods—DESE, S-LDSC, and scDRS—across 19 complex traits. Our analysis reveals a fundamental trade-off between detection sensitivity and biological specificity. We demonstrate that while S-LDSC identifies extensive spatial signals, it suffers from inflated non-specific significant associations. Conversely, scDRS is highly specific but conservative, performing well only in tissues with strong biological signals while missing subtle associations in sparser datasets. DESE overcomes these limitations, consistently achieving high power and robust specificity across both simulated and real-world scenarios. SMECT provides critical guidelines for method selection and serves as a foundational resource for developing robust spatial analyses of human complex traits. The framework is publicly available at https://github.com/pmglab/smect.

## Introduction

Genome-wide association studies (GWAS) have successfully identified thousands of genetic loci associated with complex human traits and diseases^1–3^. However, translating these statistical signals into actionable biological insights remains a major bottleneck, primarily because the specific cell types and spatial contexts in which these risk variants manifest are often unknown^4–6^. Spatially resolving the cellular basis of complex traits is therefore critical for elucidating disease mechanisms and prioritizing therapeutic targets. In psychiatric disorders like schizophrenia, for instance, pinpointing the specific brain regions or neuronal subtypes where genetic risk variants exert their effects is essential for developing targeted interventions^7–9^.

To bridge this gap, a burgeoning class of post-GWAS computational methods has emerged, aiming to integrate genetic association data with spatial transcriptomics. While these methods share the common hypothesis that trait-associated genes exhibit distinct spatial expression patterns^7,10–12^, they differ fundamentally in their statistical frameworks— ranging from heritability partitioning^10^, polygenic expression^11^, to iterative gene-set enrichment^12^. Despite their increasing application, a critical gap remains: the absence of a systematic, unbiased evaluation of their comparative performance. Given the inherent sparsity, high noise levels, and complex spatial dependencies of current spatial transcriptomics technologies, the reliability and reproducibility of these tools are largely unknown. This raises a fundamental question: to what extent can we trust the spatial mappings of complex traits currently reported in the literature?

Here, we address this challenge by developing SMECT (Spatial Mapping Evaluation of Complex Traits), the first comprehensive framework for benchmarking methods that integrate GWAS summary statistics with spatial transcriptomics. SMECT combines a real genotype-based phenotype simulation, a biologically realistic modeling of spatial autocorrelation, and technical noise simulation, with a curated collection of 21 diverse real-world datasets and a multi-faceted assessment toolkit. Using this framework, we systematically evaluated three state-of-the-art methods—DESE, S-LDSC, and scDRS— across 19 complex diseases and traits.

## Results

### Overview of SMECT

To address the critical need for standardized evaluation of spatial mapping tools, we developed SMECT (Spatial Mapping Evaluation of Complex Traits), a comprehensive benchmarking framework composed of three integrated modules designed to stress-test method performance across diverse biological scenarios (Fig. 1a).

**Fig. 1:**
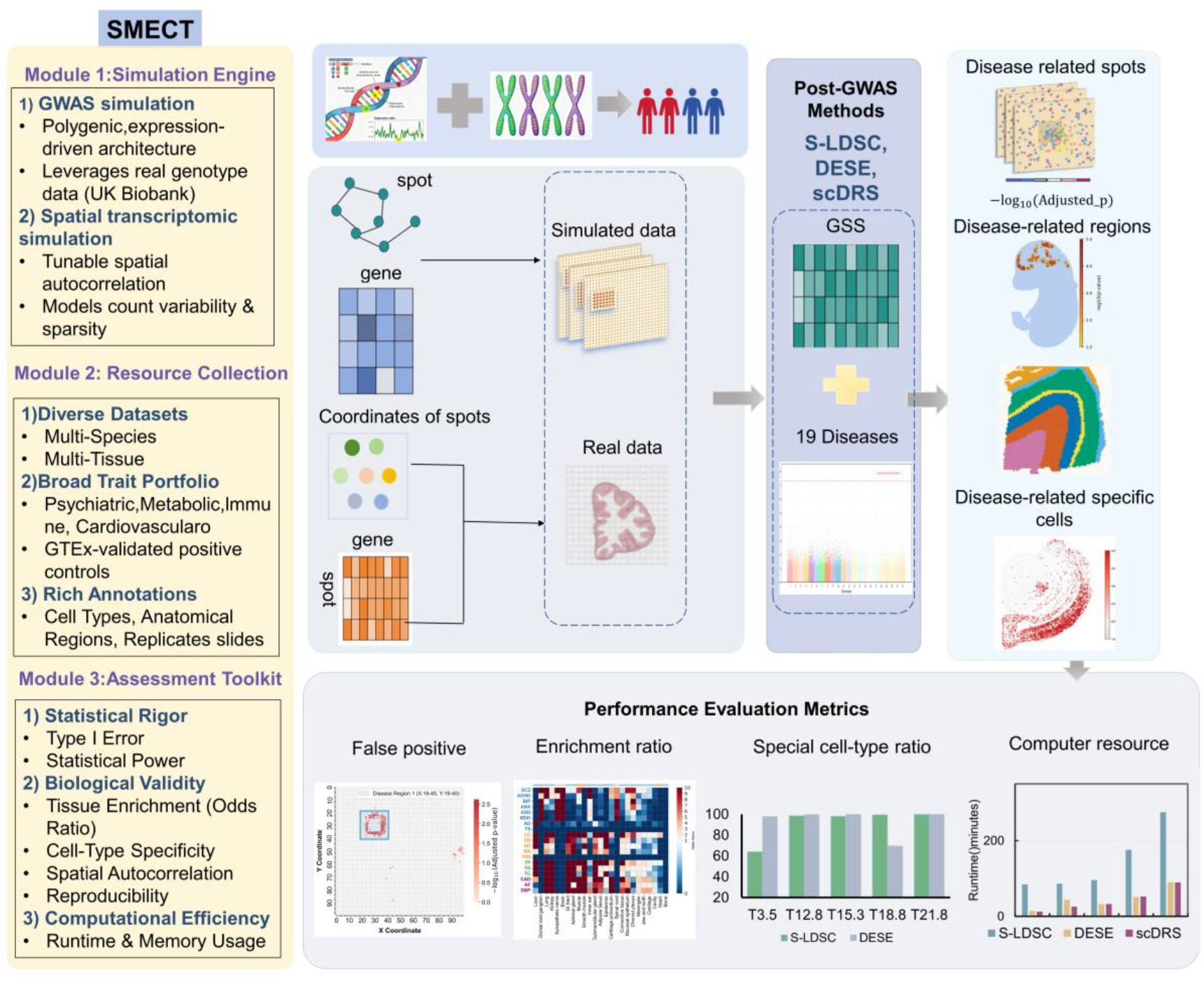
Benchmarking workflow of SMECT. Schematic overview of the SMECT benchmarking workflow, illustrating its three-module structure: Simulation Engine, Resource Collection, and Assessment Toolkit. The workflow processes real and simulated spatial transcriptomics (ST) data to evaluate three post-GWAS mapping methods. Performance is assessed across simulations, cell-type specificity, and computational resources.

#### Module 1: Simulation engine for ground-truth benchmarking

Evaluating statistical validity requires ground-truth data that real experiments cannot provide. To address this, SMECT establishes a rigorous simulation pipeline to generate synthetic datasets with known causal architectures. The engine first simulates realistic GWAS summary statistics using a hierarchical model where phenotype heritability is mediated by gene expression, leveraging real genotype data from the UK Biobank. Complementing this, it generates high-fidelity spatial transcriptomics data with explicit control over key confounding factors, including spatial autocorrelation (via a Matérn covariance kernel^13^), data sparsity (via tunable dropout rates), and count overdispersion. Crucially, the simulation allows for the designation of disease susceptibility genes as spatially variable features, modeling their localized enrichment within predefined phenotypic regions. This controlled system enables precise quantification of Type I error (null hypothesis) and, separately, of specificity and statistical power (alternative hypothesis).

#### Module 2: A curated, multi-species resource collection

To evaluate method generalizability, we curated 21 real-world spatial transcriptomics datasets covering three species (human, macaque, and mouse) and diverse technological platforms (including Stereo-seq^14^, 10x Genomics Visium^15^, and STARmap^16^) (Fig. 1b).

We integrated these resources with GWAS summary statistics for 19 complex traits spanning psychiatric, cardiovascular, immune, and metabolic domains. To validate the quality of the genetic input, we performed a positive control analysis using bulk RNA-seq data from the Genotype-Tissue Expression (GTEx) project. This confirmed that the selected GWAS statistics retain sufficient signal to recover known tissue-specific associations (Extended Data Fig. 1), ensuring that any potential lack of signal in downstream analyses stems from the spatial mapping methods or the spatial data itself, rather than deficiencies in the genetic input.

#### Module 3: Multi-Faceted Assessment Toolkit

SMECT employs a multi-dimensional evaluation pipeline to quantify performance beyond simple accuracy. We assess statistical rigor (Type I error and power), biological validity (tissue-level enrichment odds ratios and cell-type specificity), and spatial coherence (Moran’s I of genetic enrichment signals). Additionally, the toolkit evaluates reproducibility across technical replicates and quantifies computational efficiency (runtime and memory usage), providing a holistic view of each method’s practical utility.

### Performance evaluation using simulated data

To rigorously evaluate the performance of S-LDSC, DESE, and scDRS in the absence of a ground-truth standard for disease–cell localization in real data, we first applied these methods to simulated data generated by the SMECT pipeline. In each simulation scenario, we generated 100 sets of 100 × 100 spatial grids containing 20,000 genes. In short, in the simulated high-resolution dataset, each grid represents a cell, similar to the spatial transcriptome data generated by Stereo-seq.

We first established their baseline statistical behavior by evaluating the control over type I errors of the three methods and the specificity. In null simulations (those lacking any disease-associated spots in a slice), all three methods maintained a false positive rate (FPR) of zero. However, performance diverged under the alternative hypothesis, where we introduced a true disease signal and assessed “spillover” effects—non-specific detection outside the defined disease region in the same slices. While scDRS remained highly conservative with negligible non-specific detection (1-15 spots), S-LDSC and DESE exhibited distinct error profiles (Table 2). In scenarios with regional disease spots (regions 25-34), DESE produced moderate off-target signals (11 spots). In contrast, S-LDSC showed nearly double the rate of non-specific detection (21 spots). Critically, the spatial distribution of these errors differed: S-LDSC’s non-specific spots were distributed further from the true disease center compared to DESE (Fig. 2). This “signal leakage” suggests that S-LDSC’s reliance on LD scores and broad genomic annotations makes it prone to identifying spatially correlated but non-causal regions. This limitation likely drives the broad, non-specific signals observed in real datasets.

**Table 1.** Methodological comparison of three approaches for spatial mapping.

| Feature | S-LDSC | DESE | scDRS |
| --- | --- | --- | --- |
| **Core Principle** | **Partitioning Heritability.** Assumes trait heritability ($h_g^2$) is partitioned across genomic annotations. It tests if the GSS annotation for a spot explains a significant fraction of $h_g^2$. | **Iterative Enrichment of Selective Expression.** Assumes that a refined set of causal genes will show strong selective expression in relevant tissues. It employs an iterative loop to simultaneously purify the disease gene set and identify enriched spots. | **Aggregate Polygenic Expression.** Assumes that disease-relevant cells exhibit a higher cumulative expression score across a fixed set of putative disease genes. |
| **Signal Processing & Refinement** | **Static SNP-level Model.** Uses the raw GWAS $\chi^2$ statistics directly. The GSS annotation is treated as a fixed feature of SNPs to explain their aggregate genetic effect. There is no refinement of the underlying genetic signal. | **Dynamic Gene-set Refinement.** The iterative framework is its core strength. It actively uses the GSS enrichment signal to guide multiple rounds of a conditional gene-based test (by ECS), systematically **pruning indirectly associated genes**. This process distills a noisy, broad GWAS signal into a refined, more causal-like gene set for final evaluation. | **Static Gene-level Weighting.** Uses a fixed set of putative disease genes (from MAGMA). It weights each gene's GSS by its pre-computed GWAS z-score, but does not attempt to refine this initial gene list or remove potentially spurious associations within it. |
| **Statistical Test & Significance** | **Parametric Regression.** A linear regression of SNP $\chi^2$ statistics on LD scores. Significance is derived from the regression coefficient ($\tau_c$), which is sensitive to model assumptions (e.g., infinitesimal genetic architecture). | **Non-parametric Rank Test & Global Permutation.** 1. Within each iteration, a **Mann-Whitney U test** directly compares the GSS ranks of disease vs. background genes, offering a gene-level, distribution-free assessment. 2. Final p-values are derived from a **global permutation** of GSS values, which empirically corrects for selection bias from the iterative process and addresses multiple testing. | **Monte Carlo Simulation.** A weighted sum of GSS values is compared against an empirical null distribution generated from randomly sampled control gene sets. Significance depends on the quality of matching between control and disease gene sets (e.g., in expression level, variance). |
| **Hypothesized Drivers of Performance** | **Strength:**<br>Rigorously models LD.<br>**Limitation:** Its power can be limited by relying on a fixed genetic model and being an indirect test of gene function (it models SNP heritability, not gene ranks). | **Strength:**<br>Signal Purity: The iterative refinement actively removes noise from indirect genetic associations, likely boosting statistical power.<br>Robust Test: The non-parametric U-test is robust to GSS outliers and distributional assumptions, enhancing specificity.<br>Rigorous Error Control: The global permutation strategy provides a robust control of Type I error. | **Strength:**<br>True single-cell resolution and an elegant weighting scheme.<br>**Limitation:** Its performance is contingent on the initial, unrefined gene set from MAGMA. It may be less powerful if this set contains many non-causal genes, a problem DESE's iterative process is designed to solve. |
| **Output in this Benchmark** | A p-value for each individual spot. | A p-value for each individual spot. | A p-value for each individual spot. |

**Table 2:**
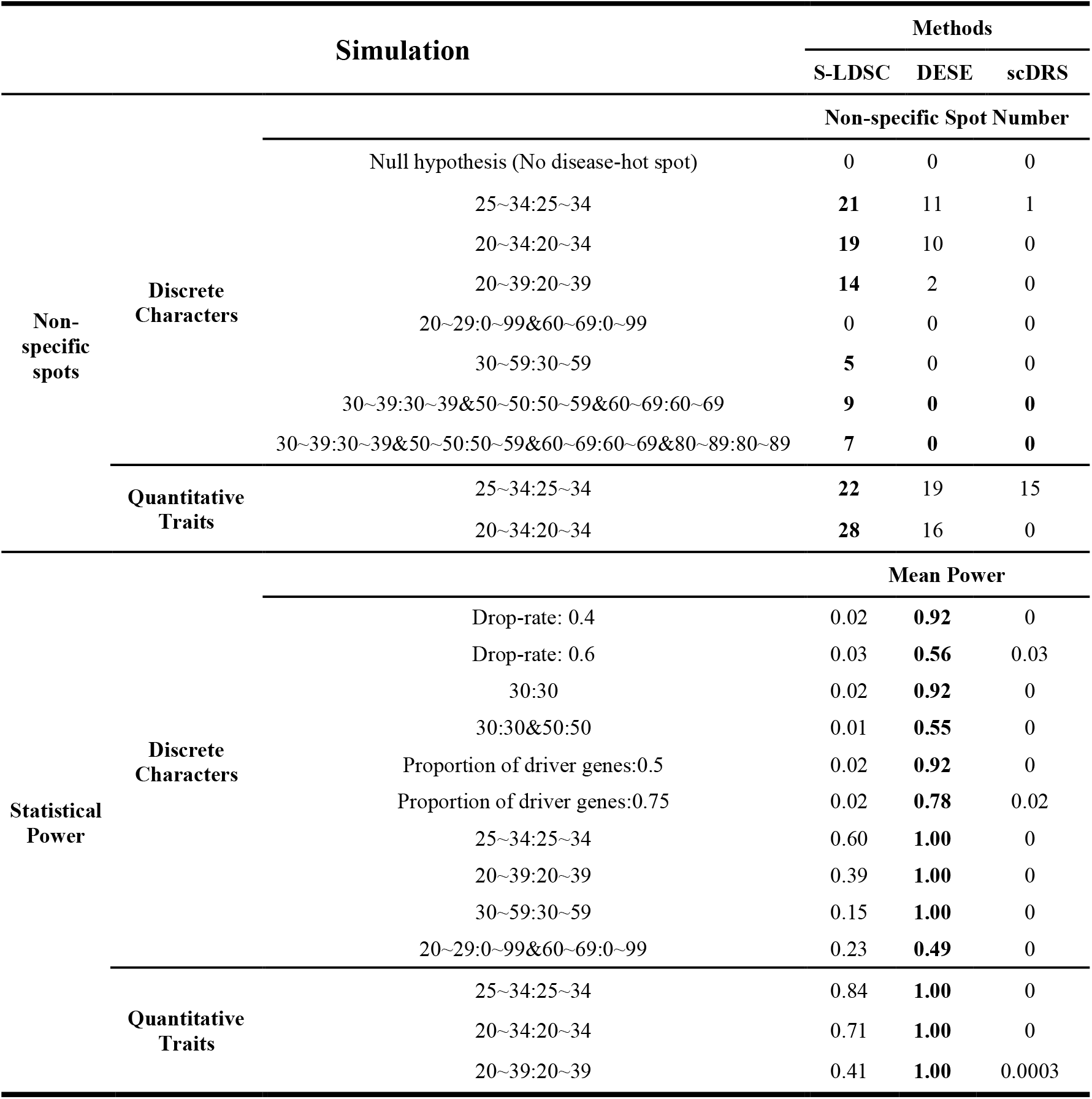
The number of non-specific spots and the power of different methods in different simulation scenarios.

**Fig. 2:**
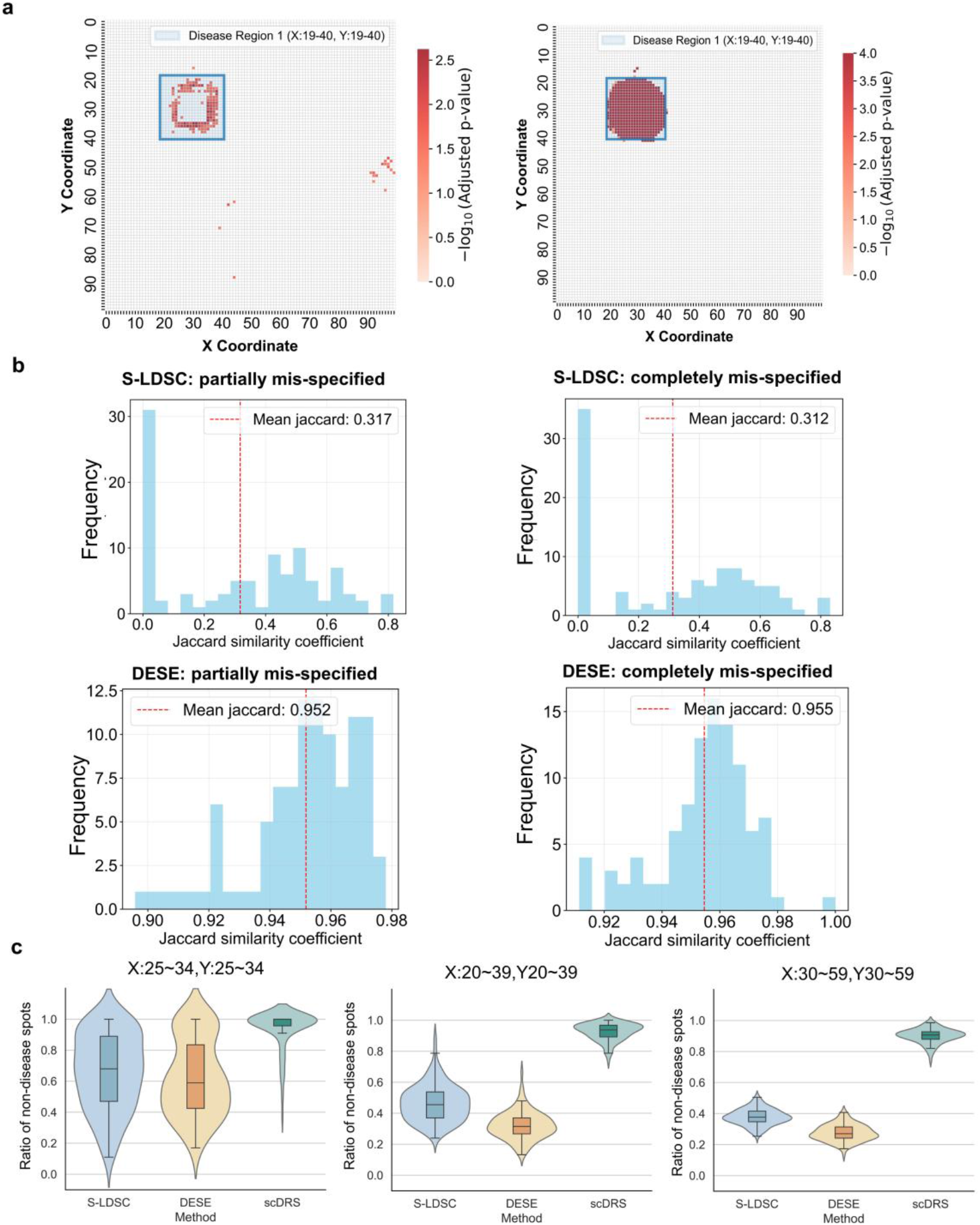
Assessment of the specificity of S-LDSC, DESE, and scDRS and robustness to mis-specified spatial annotations. **a**, Non-specific control under alternative hypothesis simulations. Non-specific spots are calculated as the number of significant spots outside the disease area (FDR < 0.05). Heatmaps of disease region 1 (X: 19-40, Y: 19-40) for DESE (left) and S-LDSC (right), showing the spatial pattern of significance, with the blue square marking the simulated “true disease region”. **b**, Robustness to mis-specified spatial annotations. Comparison of DESE and S-LDSC under three label perturbation scenarios: correct labels, partially mis-specified labels, and completely mis-specified labels. Corresponding histograms of the Jaccard similarity coefficient between the inferred and simulated “true disease regions” for partially and completely mis-specified labels. Red dashed lines indicate the mean Jaccard index. **c**, Violin plots with overlaid boxplots displaying the ratio of non-simulated disease hotspots among the top 100, 400, and 900 spots ranked by z-score (without multiple testing correction), as detected by three methods (S-LDSC, DESE, scDRS). The three panels correspond to analyses across distinct coordinate ranges: left panel (X: 25–34, Y: 25–34), middle panel (X: 20–39, Y: 20–39), and right panel (X: 30–59, Y: 30–59). Detailed information for the violin plots is presented in Supplementary Table 5-7.

Next, we evaluated statistical power under the alternative hypothesis. We first modeled a challenging scenario: a single disease hotspot at spatial coordinates (30, 30) with varying levels of technical noise (dropout rates of 0.4 and 0.6). DESE demonstrated high sensitivity, maintaining power between 0.92 and 0.56 even as data sparsity increased. Conversely, both S-LDSC and scDRS failed to detect these focal signals effectively (power < 0.1 for S-LDSC; power around 0 for scDRS) (Table 2). For S-LDSC, the failure likely stems from its resolution limit; it is designed to detect heritability enrichment across broader annotations rather than pinpoint single-spot variations.

This hypothesis was confirmed when we expanded the simulation to regional hotspots (mimicking tissue-level associations). As the disease signal footprint increased, S-LDSC’s mean power improved significantly (rising to 0.60–0.84), though it still lagged behind DESE, which achieved near-perfect power (0.99–1.00) (Table 2). Notably, scDRS almost did not detect significantly correlated signals in the simulated data, showing the minimum power. Nevertheless, a closer inspection of z-score rankings prior to multiple testing correction showed that subsets of slices harboring a higher proportion of simulated disease hotspots remained among the top 100 spots (the highest proportion of disease spots in the top 100 rankings among the 100 simulated slices, = 0.63; Supplementary Table 7). The distribution of power in the 100 slices of S-LDSC varies greatly (Extended Data Fig. S3). These results clarify the operational boundaries of the methods: DESE is sensitive to both focal and regional patterns, whereas S-LDSC requires broader spatial signals to achieve statistical significance. We also simulated multi-region hotspot scenarios under alternative assumptions (Extended Data Fig. 4a). DESE still shows high mean power (0.67-1). However, the average power of S-LDSC remains relatively low (0.08-0.26), and it exhibits high efficacy in only a small number of slices across different simulation scenarios (Extended Data Fig. 4b).

To evaluate specificity according to the ranking instead of statistical p-values, we sorted the results (before multiple testing correction) by z-scores in descending order and, for three simulation scenarios initialized with 100, 400, and 900 disease spots, respectively, calculated the ratio of disease spots among the top 100, 400, and 900 spots in each scenario. Across all scenarios, scDRS identified the highest ratio of non-disease hotspots, reflecting poor specificity (mean ratio: 0.94, 0.93, 0.90; Fig. 2c). S-LDSC identified a higher proportion of non-disease hotspots compared with DESE, indicating that S-LDSC has lower specificity. Notably, DESE exhibited the highest specificity among all the methods tested, with respective mean ratio values of 0.62, 0.32, and 0.28, versus 0.66, 0.46, and 0.38 for S-LDSC.

To evaluate the robustness of the methods to incorrect spatial labels when generating the gene specificity scores (GSS) based on the transcriptomic spatial data, we perturbed the spatial labels in the preprocessing by the gsMap^7^. Specifically, we introduced errors into the spot annotations within the disease region before GSS calculation, creating scenarios where the input features were derived from partially or completely mis-specified labels. We then assessed how well each method could recover the true disease locations despite these corrupted input scores. To quantitatively evaluate the overlap between the predicted disease regions and the experimentally validated ground-truth regions, we employed the Jaccard index — a widely adopted metric for measuring the similarity of two sets. DESE exhibited remarkable stability, maintaining high concordance with the simulated “true disease region” (a Jaccard index value of around 0.95) even when the underlying GSS was based on noisy labels. In contrast, the performance of S-LDSC deteriorated significantly with a Jaccard index value of around 0.31 (Fig. 2b). This disparity highlights a practical advantage of DESE: its iterative refinement approach allows it to correct for initial labeling errors by homing in on the underlying expression signal, whereas the quality of the predefined input annotations constrains S-LDSC.

While our SMECT simulation pipeline provides a comprehensive evaluation, a limitation should be acknowledged. The simulation, although designed to capture key biological parameters, relies on simplified assumptions—such as predefined circular disease regions and Gaussian decay functions—that may not fully reflect the intricate spatial patterns in real biological tissues. Therefore, we also performed extensive testing on real-world datasets.

### Accuracy of methods in spatial mapping of complex traits

To ensure the validity of our genetic inputs, we first performed a positive control analysis using bulk RNA-seq data from the Genotype-Tissue Expression (GTEx) project^17^. This step robustly replicated canonical trait–tissue associations, including the link between schizophrenia and the frontal cortex (BA9)^18,19^ and between total cholesterol and the liver^20–22^(Extended Data Fig. 1). With the biological signal of the GWAS datasets confirmed in bulk tissue, we proceeded to evaluate method performance on the more complex, high-sparsity regimes of spatial transcriptomics.

To validate the methods on complex biological data, we analyzed a spatial transcriptomics dataset of a mouse embryo at day 16.5 (data labeled E16.5), which captures 25 distinct tissues (Fig. 3a). We leveraged GWAS summary statistics for 19 human traits, capitalizing on the high conservation of gene expression profiles between mouse and human. Method performance was assessed by computing odds ratios (*ORs*) for each tissue region to indicate enrichment of significant tissues (See more in Method).

**Fig. 3:**
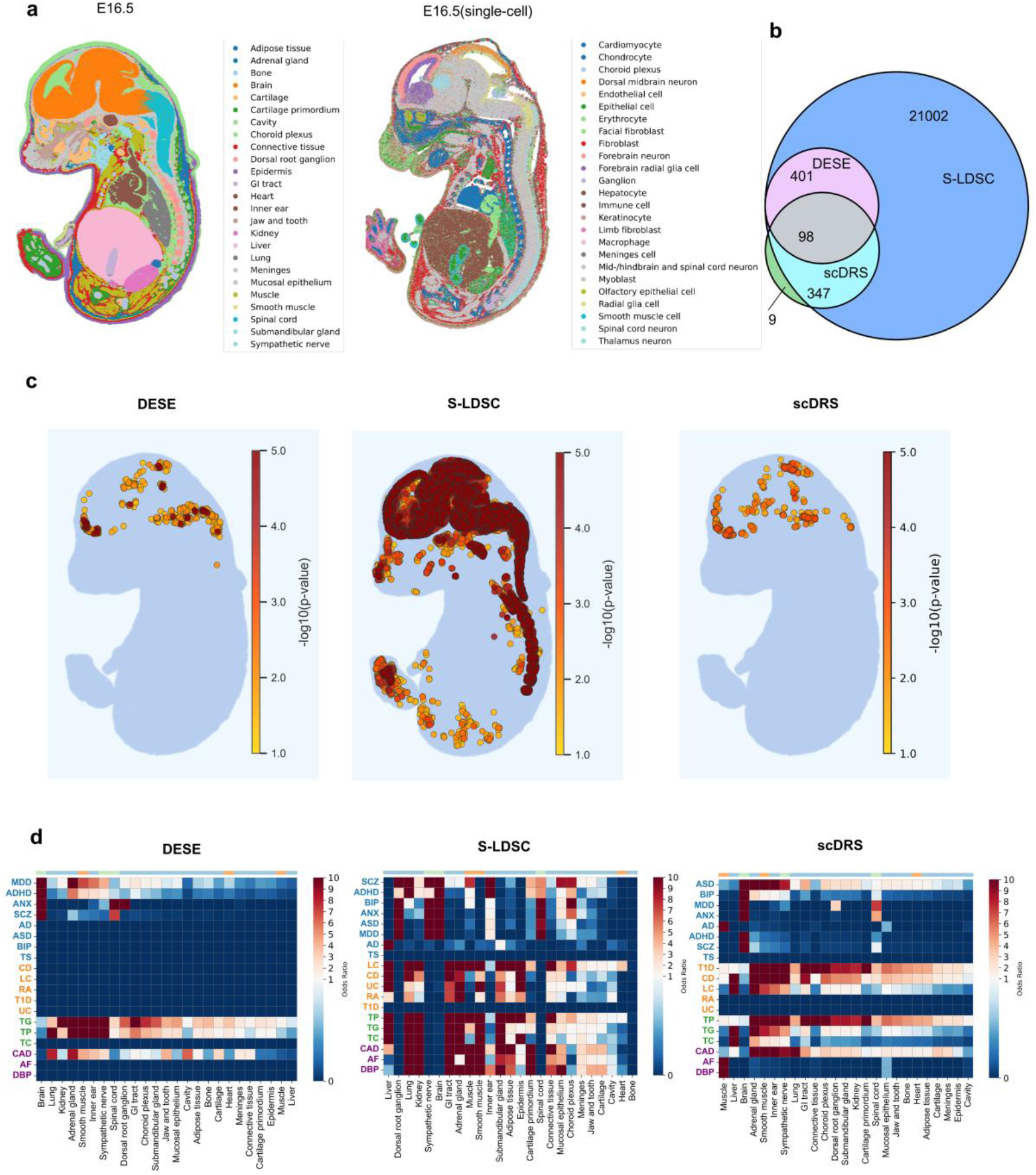
Validation of methods using a mouse embryo E16.5 spatial transcriptomics dataset. **a**, Spatial mapping of tissue types (left) and single-cell type annotations (right) in a mouse embryo at embryonic day 16.5 (E16.5). **b**, Venn diagram illustrating the overlap of significant trait-associated spots for schizophrenia (SCZ) identified by DESE, S-LDSC, and scDRS. **c**, Spatial distribution of significant spots for SCZ enrichment detected by DESE (left), S-LDSC (middle), and scDRS (right). Color scale indicates the negative log10 (adjusted p-value). **d**, Heatmaps showing enrichment results for 19 human traits across various tissues for DESE (left), S-LDSC (middle), and scDRS (right). Detailed information for the heatmaps is presented in Supplementary Table 1-3.

We first focused on schizophrenia (SCZ), a disorder with well-established brain-centered pathophysiology. All three methods correctly identified the brain as a significantly trait-associated tissue, yet their enrichment patterns differed substantially. S-LDSC identified numerous significant tissue-specific associations (Fig. 3b); however, these associations were broadly distributed across both neural and non-neural tissues, leading to biologically implausible associations in non-neural regions such as the cartilage primordium (*OR* = 4.21; Fig. 3c). In contrast, DESE and scDRS demonstrated greater precision and specificity. DESE identified 401 significant signals (Fig. 3b), almost exclusively concentrated in the central nervous system (CNS), including the brain and spinal cord (Fig. 3c). Similarly, scDRS identified 445 significant signals primarily within the CNS, with minimal background noise in other tissues (Fig. 3b). Notably, 98 significant signals were shared between the DESE and scDRS results. To evaluate the specificity of each method, we sorted the results by odds ratio (OR) values, where an OR > 1 indicates a positive association with SCZ susceptibility. In the DESE results, only the brain and spinal cord showed OR values greater than 1 (brain: *OR* = 22.69; spinal cord: *OR* = 7.25; Supplementary Table 1). For scDRS, only the brain exhibited an OR value > 1 (*OR* = 174.76; Supplementary Table 3). In contrast, the top five tissues with the highest OR values in the S-LDSC results were the dorsal root ganglion (*OR* = 4092.54), sympathetic nerve (*OR* = 1238.10), inner ear (*OR* = 890.80), brain (*OR* = 546.90), and lung (*OR* = 14.23; Supplementary Table 2). These results indicate that DESE and scDRS have high specificity, while S-LDSC has lower specificity, which is consistent with our previous simulation results.

This trade-off between sensitivity and specificity was consistent across the 19 traits analyzed (Fig. 3d). For example, DESE identified highly specific associations, such as robust enrichment in the brain for major depressive disorder (*OR* = 270.7). scDRS also identified highly specific associations, including a strong enrichment of total cholesterol in the liver (*OR* = 1076.2). S-LDSC, in contrast, consistently identified a broader range of associations, many of which implicated tissues lacking a clear biological connection to the trait. Examples include coronary artery disease (CAD, cartilage primordium: *OR* = 22.0), diastolic blood pressure (DBP, cartilage primordium: *OR* = 14.7), and atrial fibrillation (AF, cartilage primordium: *OR* =12.1), for which there is currently no established association between cardiovascular disease and cartilage primordium. These real-world data underscore a fundamental trade-off between sensitivity and specificity inherent in these methods.

### Cell-type specificity in single-cell resolution datasets

Single-cell resolution spatial transcriptomics enables direct assessment of cell type-specific disease associations. We analyzed a dataset from five coronal slices of the macaque claustrum^24^, a region implicated in psychiatric disorders^25^ (Fig. 4a). Given the lack of significant findings from scDRS in this dataset, we focused our comparison on DESE and S-LDSC.

**Fig. 4:**
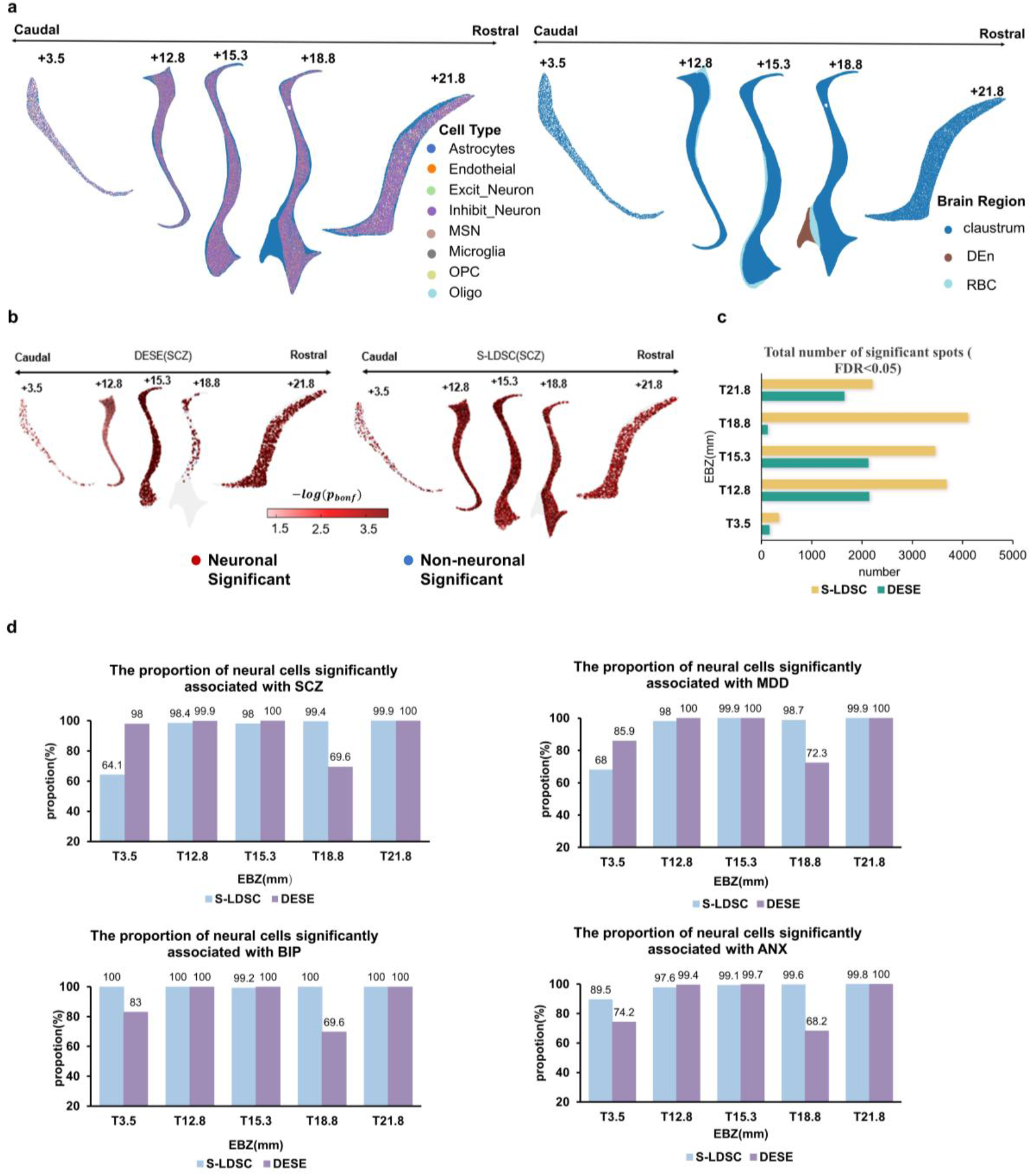
Cell-type specific disease associations in the macaque claustrum. **a**, Spatial distribution of cell types (left) and brain regions (right) across five coronal slices of the macaque claustrum. **b**, Spatial distribution of schizophrenia (SCZ)-associated cells detected by DESE and S-LDSC. Significant spots (FDR < 0.05) are colored by p-value intensity. Neuron-related and non-neuronal significant cells are marked with blue and red circles, respectively. **c**, The number of significant disorders, including major depressive disorder (MDD), associated with DESE and S-LDSC. **d**, Proportion of neural cells significantly associated with various psychiatric disorders, comparing DESE and S-LDSC across the five slices.

Given the established direct association between neuron-related cells and the development of psychiatric disorders, our analysis primarily focused on this cell population^26–28^. Both DESE and S-LDSC effectively identified enrichment in the macaque brain. S-LDSC consistently identified a substantially larger number of disease-associated cells across all slices (Fig. 4c). For instance, at the +18.8 mm position, S-LDSC detected 4,103 significant spots for SCZ, whereas DESE identified only 115. Conversely, DESE exhibited superior specificity for neuron-related cells, the primary cell population implicated in psychiatric disorders. For SCZ, DESE consistently attributed a higher proportion of significant associations to neurons compared to S-LDSC across most slices (e.g., 98% vs. 64.1% at the +3.5 mm position) (Fig. 4d). This trend was also observed for major depressive disorder (MDD). These findings highlight a nuanced trade-off: S-LDSC offers greater sensitivity in detecting associated cells, while DESE provides superior specificity in pinpointing the most biologically relevant cell types. Given the central role of the brain in psychiatric disorders, we extended our analyses to a single-cell resolution spatial transcriptomic dataset of the mouse brain^29,30^ (Fig. 5a). Prior literature has highlighted glutamatergic (Glu) neurons as a key cell type most directly implicated in the pathophysiology of psychiatric disorders, including major depressive disorder (MDD) and schizophrenia (SCZ)^31,32^. Consistent with prior observations, scDRS failed to yield significant associations and was therefore excluded from further analyses. Aligning with previous findings, both DESE and S-LDSC detected disease-associated spatial regions. Notably, S-LDSC identified substantially more significant spots than DESE (e.g., 8359 vs. 3496 spots for MDD), reflecting greater sensitivity (Fig. 5b,c). Nevertheless, DESE exhibited superior cell-type specificity. Across disorders, including SCZ, anxiety disorder (ANX), and bipolar disorder (BIP), DESE consistently detected a higher proportion of significant glutamatergic (Glu) neuron-containing spots relative to S-LDSC (e.g., 100% vs. 83% for SCZ) (Fig. 5d). This confirms that DESE is better suited to pinpointing specific cell types critical to disease pathophysiology. In contrast, S-LDSC provides a more comprehensive and sensitive characterization of potentially associated cells.

**Fig. 5:**
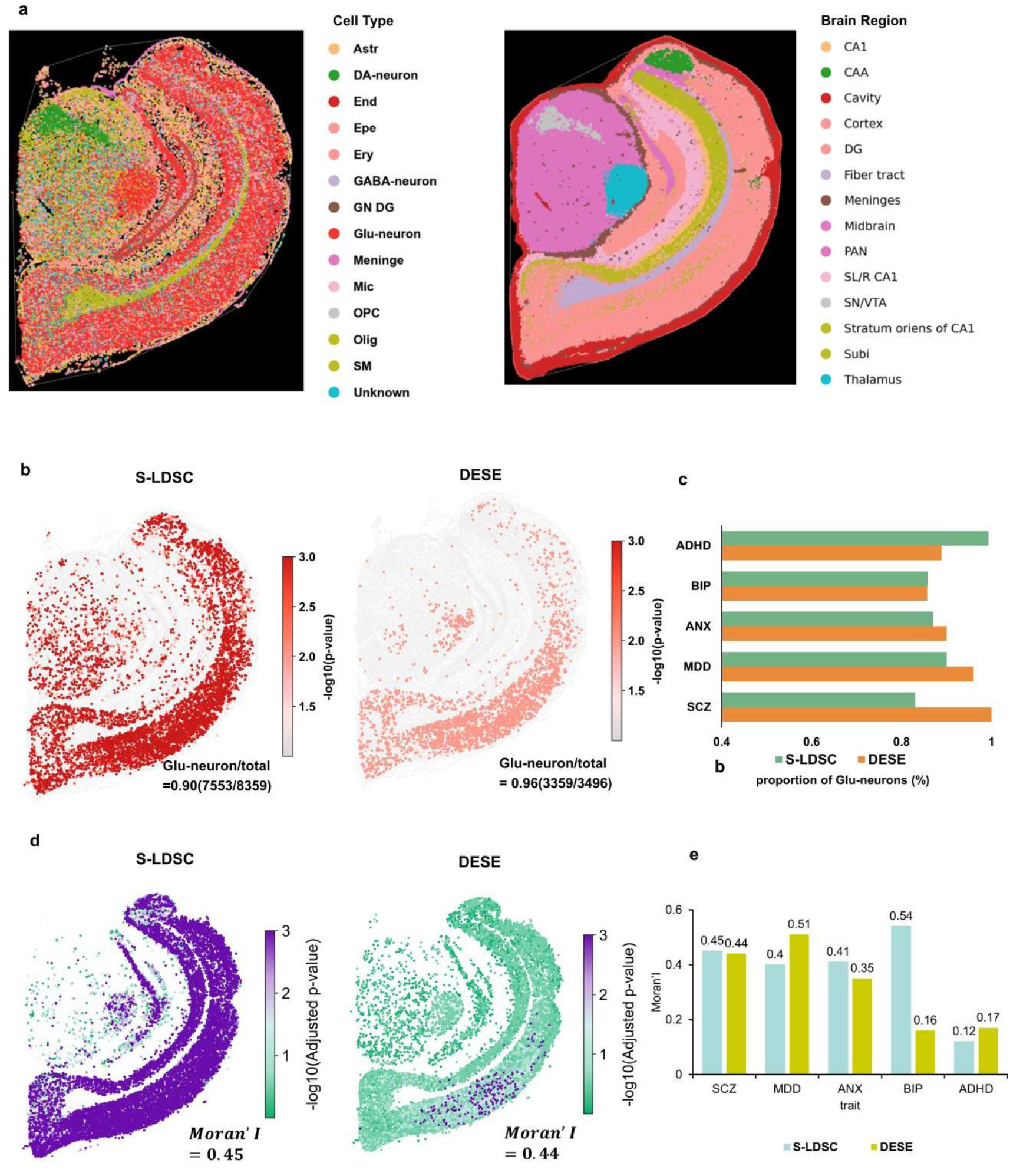
Integrative analysis of cell-type-specific and spatial disease associations in the adult mouse brain. **a**, Spatial distribution of cell types (left) and brain regions (right) in an adult mouse brain section. **b**, Spatial distribution of significant spots for schizophrenia (SCZ) detected by S-LDSC (left) and DESE (right). Red dots indicate significant spots (FDR < 0.05). **c**, Proportion of significant glutamatergic (Glu-neuron) cells associated with various psychiatric disorders for S-LDSC and DESE. **d**, Spatial distribution of p-values for schizophrenia (SCZ) association for S-LDSC (left) and DESE (right). Color scale indicates the negative log10 (adjusted p-value). **e**, Moran’s I value quantifying the spatial autocorrelation of p-values within Glu-neurons for various psychiatric traits, comparing S-LDSC (blue) and DESE (yellow).

We explored the spatial organization of genetic signals by performing Moran’s I analysis on the p-values within Glu-neurons in the adult mouse brain dataset. A higher Moran’s I value indicates stronger spatial autocorrelation. Spatial autocorrelation refers to the absence of a completely random spatial distribution, instead reflecting a tendency for similar or correlated patterns to occur in spatially proximate regions. Literature reports indicate that gene expression exhibits spatial specificity: the same cell type displays distinct expression profiles across different spatial locations, resulting in variable degrees of association with diseases^33–35^. Both S-LDSC and DESE results showed significant spatial autocorrelation for psychiatric disorders (Fig. 5d). S-LDSC identified moderate to strong spatial autocorrelation across multiple disorders, with the highest value for BIP (*Moran’s I* = 0.54, *p* < 0.05), indicating that associated Glu-neurons are spatially clustered (Fig. 5e). In contrast, DESE’s results were more trait-dependent. It revealed strong spatial clustering for MDD (Moran’s I = 0.51, *p* < 0.05) but a considerably weaker pattern for BIP (*Moran’s I* = 0.16, *p* < 0.05). These findings suggest the methods capture different facets of the spatial architecture of disease risk, with S-LDSC identifying broader correlated regions and DESE pinpointing more trait-specific clustering patterns.

### DESE and S-LDSC show robust predictions across replicate tissues

To evaluate the stability of predictions, we analyzed four adjacent spatial transcriptome slices from the dorsolateral prefrontal cortex (DLPFC) of a single human donor. As scDRS generated no significant enrichment signals, we focused on S-LDSC and DESE. Given that each spot was annotated with its corresponding cortical layer, we then aggregated these p-values to derive a layer-specific p-value for each slice using the Cauchy combination test. We calculated layer-specific p-values for SCZ association across all four slices and then computed Spearman’s rank correlation coefficients for all pairwise combinations. Both methods demonstrated robust and stable performance, yielding highly consistent disease association profiles. DESE exhibited remarkable consistency, with pairwise correlation coefficients ranging from 0.67 to 0.88 (Fig. 6). S-LDSC also showed good stability, with correlations spanning 0.54 to 0.86. The high consistency observed for both methods affirms their reliability for generating stable, spatially resolved genetic association analyses.

**Fig. 6:**
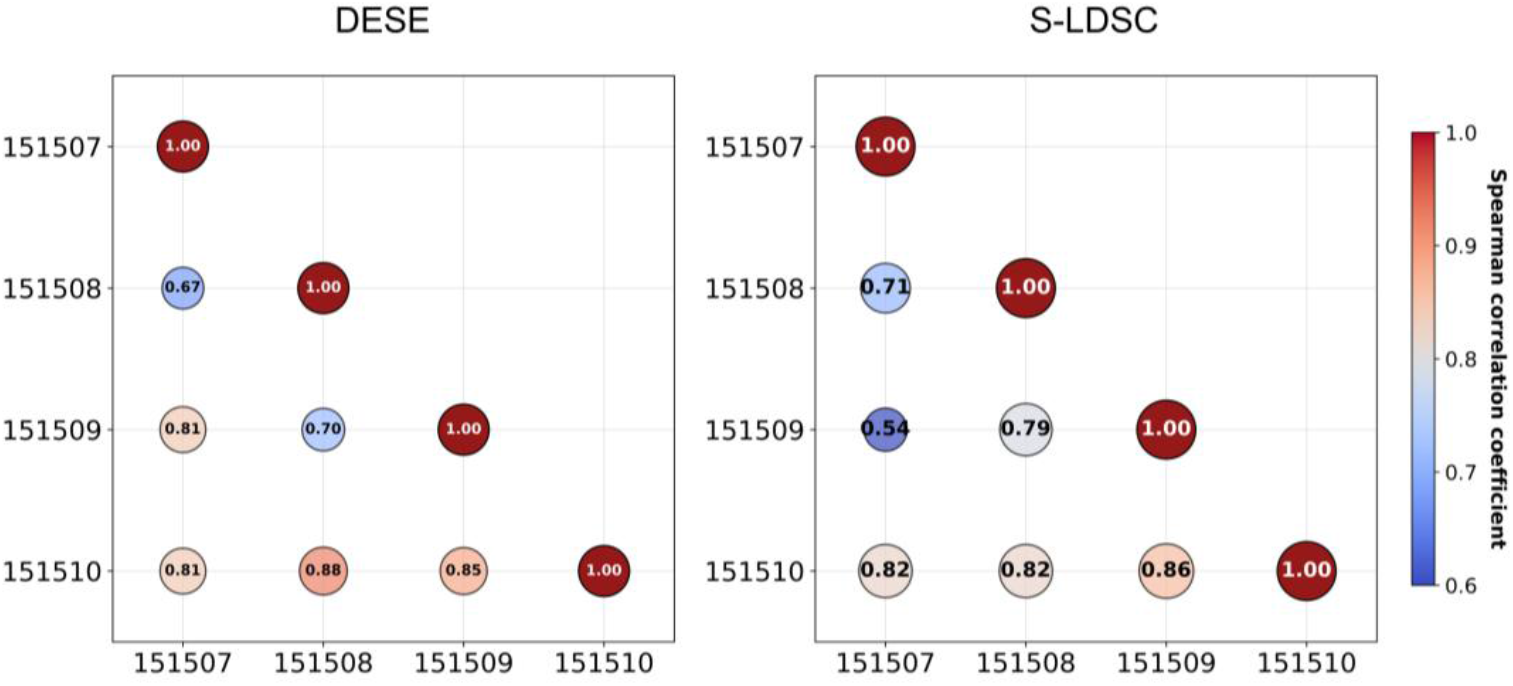
Robustness of trait associations across replicate human DLPFC slices. Pairwise Spearman’s rank correlation coefficients of layer-specific p-values for schizophrenia (SCZ) association across four replicate slices for DESE (left) and S-LDSC (right). The color and numerical value in each circle indicate the strength of the correlation.

### Computational performance

Finally, we benchmarked the computational resource consumption of the three methods. To ensure a fair comparison, all methods were executed in single-thread mode. S-LDSC consistently required the longest runtimes (e.g., 275.20 minutes for the mouse embryo dataset, compared to 90.70 min and 90.10 min for DESE and scDRS, respectively). Conversely, DESE incurred the highest memory usage (53.90 GB) compared to S-LDSC (34.90 GB) and scDRS (26.80 GB) (Fig. 7a, b). While scDRS was generally the most resource-efficient, DESE benefits significantly from multi-threading capabilities, allowing for substantial acceleration on multi-core systems. For instance, utilizing 8 threads reduced the runtime for the mouse embryo dataset from 153 minutes (2 threads) to 82 minutes (Fig. 7c). This highlights a practical trade-off: while scDRS offers the smallest resource footprint, DESE achieves superior speed via parallel processing when sufficient memory is available.

**Fig. 7:**
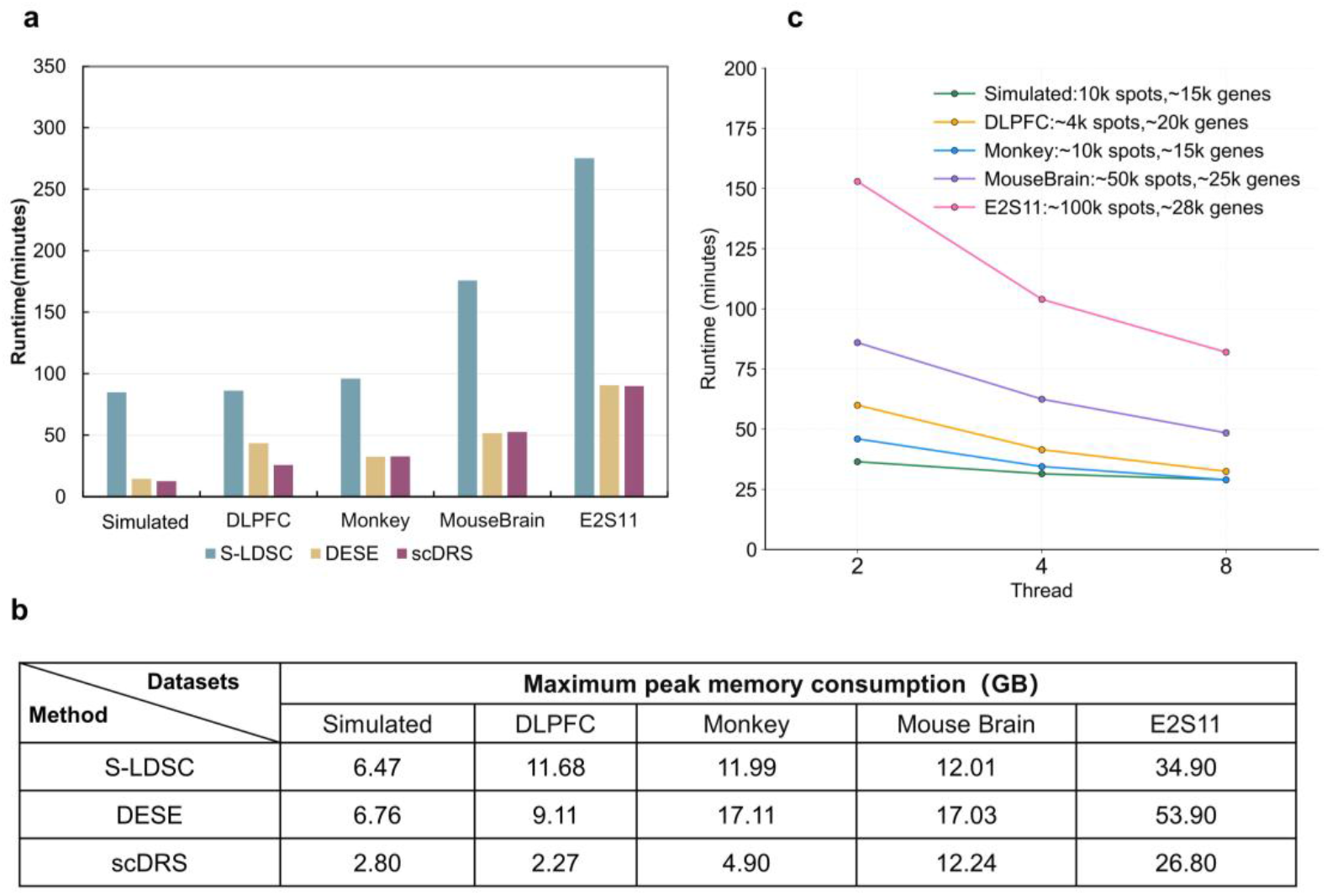
Computational performance of S-LDSC, DESE, and scDRS. **a**, Runtime comparison (in minutes) on various simulated and real-world datasets. **b**, Peak memory usage (in GB) for S-LDSC, DESE, and scDRS across the same datasets. **c**, Impact of multi-threading on DESE runtime as a function of the number of threads used.

## Discussion

We established SMECT to rigorously evaluate post-GWAS methods for mapping cell types associated with human complex traits. By integrating real genotype-based simulations and 21 empirical datasets into SMECT, our analysis uncovers a defining trade-off between sensitivity and specificity across DESE, S-LDSC, and scDRS that dictates their optimal application. We demonstrate that while S-LDSC offers high sensitivity, it is prone to inflated non-specific significant associations that can lead to biologically implausible associations in complex tissues. Conversely, scDRS exhibits high specificity but is conservative, showing limited power in sparse data regimes. DESE emerges as a robust alternative, effectively balancing statistical power with stringent error control. By delineating the boundaries of these methods, SMECT provides critical guidance for method selection and serves as a foundational resource for ensuring robust and interpretable spatial analyses of human complex traits.

Our simulations exposed critical disparities in Type I error control. While DESE and scDRS maintained strict error rates, S-LDSC exhibited moderately inflated non-specific significant associations. This divergence from prior reports^7^ likely stems from our simulation design: rather than randomly distributing causal variants, we explicitly modeled genomic regions with high background heritability that overlap with spatial annotations. This suggests that S-LDSC may misattribute genetic signal from confounding regulatory elements to spatial features, driving false discoveries.

Regarding statistical power, DESE proved the most sensitive, likely due to its conditional association framework that iteratively refines the causal gene set to remove indirect associations. Conversely, S-LDSC’s aggregation of gene expression into coarse genomic annotations appears to sacrifice resolution. While scDRS offers high specificity, its low sensitivity in our simulations implies a dependence on strong, pre-refined signals—a condition often unmet in sparse data. This establishes a clear spectrum: DESE balances power and precision; S-LDSC offers broad sensitivity at the cost of specificity; and scDRS prioritizes specificity but risks false negatives.

These simulation-derived profiles were corroborated by empirical analyses. In the mouse embryo, S-LDSC implicated biologically implausible tissues (e.g., cartilage) for psychiatric traits, reflecting the “signal leakage” observed in simulations. In single-cell resolution datasets (macaque and adult mouse brain), S-LDSC identified vastly more cells than DESE, yet DESE attributed a significantly higher proportion of its signals to disease-relevant neurons. This confirms DESE’s superior specificity in pinpointing driver cell types.

These findings suggest distinct use cases. S-LDSC is viable for broad, exploratory hypothesis generation, provided results are interpreted with caution regarding non-specific significant associations. For mechanistic studies requiring precise cell-type localization, DESE is the more robust choice. Practically, scDRS remains a resource-efficient option, though DESE’s parallelization effectively mitigates its computational cost. Furthermore, the ability of these methods—particularly DESE—to recapitulate human GWAS biology in cross-species spatial data validates the utility of model organisms for spatial genetics. We propose that integrating the granular specificity of DESE with the broad scope of S-LDSC could yield complementary insights.

Limitations of our study should be acknowledged. First, our simulations relied on simplified geometric assumptions; future benchmarks should incorporate data-driven spatial topologies. Additionally, current specificity metrics rely on pre-existing cell annotations. Developing methods for de novo identification of disease-relevant cell states would further refine these evaluations. Finally, the variable performance of scDRS warrants further investigation into optimal parameter tuning.

In summary, SMECT provides a rigorous standard for the evolving field of spatial trait mapping. By systematically delineating the strengths and boundaries of current tools, this framework offers essential guidance for researchers and establishes a foundation for developing more accurate and biologically interpretable computational methods.

## Methods

### Data curation and preprocessing

We curated a collection of 21 publicly available spatial transcriptomics datasets spanning three species (human, macaque, and mouse) and multiple technologies (Supplementary Data). For post-GWAS analysis, we collected summary statistics for 19 complex diseases and traits from public repositories (Supplementary Data).

A standardized pipeline was applied to all spatial transcriptomics datasets. Raw gene expression counts were log-transformed and normalized. To mitigate data sparsity, we employed the gsMap method^7^ to compute a gene-specific score (GSS) for each of the top 3,000 highly variable genes per dataset. These GSS values served as the common input for the three benchmarked post-GWAS methods.

### Gene-SNP annotation

For all methods, single nucleotide polymorphisms (SNPs) from GWAS summary statistics were assigned to a gene if they were located within its transcribed region or 10 kb flanking regions^36^. This window is a standard approach to capture potential cis-regulatory elements.

### Simulation framework

As a core component of the **SMECT** framework, we developed a comprehensive simulation pipeline to rigorously evaluate method performance. This pipeline generates biologically realistic spatial transcriptomics data with explicit control over spatial autocorrelation, sparsity, and count variability using a Matérn covariance kernel.

To generate corresponding GWAS summary statistics, we implemented a hierarchical model that simulates a quantitative phenotype mediated by gene expression, linking genetic variants (eQTLs) to gene expression levels and subsequently to the final phenotype. This approach allows for the creation of gold-standard datasets for both null (Type I error) and alternative (statistical power) hypothesis testing. The whole-genome 4,206,964 variants with minor allele frequency >0.05 of 490,541 subjects from UKB were used for the simulation in the present study. A detailed description is provided in the Supplementary Methods.

### Parameter settings

We tested the phenotypes of three methods (S-LDSC,scDRS, DESE) on both simulated and real datasets. The parameters of each method were set as described below for each program.

S-LDSC was implemented following the instructions on the gsMap website (https://yanglab.westlake.edu.cn/gsmap/document/software).

- **Gene window size:** We used a ±10 kb window around each gene to annotate SNPs, consistent with standard practice in functional enrichment analyses.
- **LD scores:** Precomputed LD scores used in real datasets were based on the 1,000 Genomes Phase 3 European reference panel. The LD reference panel used in simulated datasets was sourced from 490,541 subjects in the UK Biobank (UKB) cohort.
- **Baseline model:** The gsMap default baseline LD model was used to control for broad functional annotations. DESE. We followed the instructions on the KGGSum website: https://pmglab.top/kggsum/. DESE has been integrated into the KGGSum platform.
- **Permutation number:** The number of phenotype permutations was set to permutation-num = 100, which balances computational efficiency with stable empirical P-value estimation across phenotypes.
- **Threshold of the adjusted p-value:** We set the p-value-cutoff = 0.05.
- **Multiple testing correction method:** We set the multiple testing correction method as gene-level multiple testing correction = bhfdr.
- **Prevalence of affected individuals:** We set the prevalence to 0.01.
- **Other settings:** All additional parameters were kept at their recommended defaults to ensure comparability across traits. scDRS. We follow the tutorial on the GitHub repository of scDRS: https://martinjzhang.github.io/scDRS/.
- **Number of control gene sets:** We set n_ctrl = 1,000, the recommended upper bound for generating matched control gene sets that account for gene-specific confounders.
- **Weight:** Gene weight options. We set weight = zscore
- **Maximum number of genes for each gene set:** We set n_max = 1,000.
- **Flag_raw_count:** Raw count matrices were log-normalized and scaled per tutorial recommendations. We set Flag_raw_count = True.
- **Flag_filter_data:** We set Flag_filter_data = True to apply minimum cell count and gene filtering.

### Performance evaluation metrics

#### Type I error control and statistical power in simulations

To assess Type I error control, we quantified the false positive rate (FPR) under 100 null simulations. Non-specific was defined as the proportion of simulation replicates in which at least one spatial spot was erroneously identified as significant after Benjamini–Hochberg correction (FDR < 0.05). Under alternative simulations, we further enumerated non-specific signals as significant spots detected outside the predefined disease-associated region.

Statistical power was evaluated using 100 alternative simulations containing known disease hotspots. For scenarios with a single hotspot, power was summarized by a weighted detection score based on a Gaussian decay kernel centered at the true causal location; detections closer to the hotspot center contributed more strongly to the overall score (see Supplementary Methods for details). For simulations with multiple hotspots, power was defined as the proportion of true disease-associated hotspots that were successfully recovered. Slice-level power values exceeding 0.8 were truncated at 1 to prevent disproportionate influence of simulations with extremely high detection confidence.

#### Enrichment and specificity in real-world datasets

In the mouse embryo dataset, we computed odds ratios (ORs) to quantify the enrichment of trait-associated spots within 25 anatomically defined tissues. For single-cell resolution datasets (macaque claustrum, adult mouse brain), we assessed cell-type specificity by calculating the proportion of significant associations localizing to biologically relevant cell types (e.g., neurons for psychiatric disorders).

#### Robustness and spatial structure analyses

To evaluate reproducibility, we analyzed four replicate spatial transcriptomics slices from the human dorsolateral prefrontal cortex (DLPFC). We computed pairwise Spearman’s rank correlation coefficients of layer-specific schizophrenia association p-values across all slices. The spatial organization of genetic signals was investigated by performing a Moran’s I spatial autocorrelation analysis on the association p-values within glutamatergic neurons in the adult mouse brain dataset.

#### Computational performance

All methods were benchmarked on a high-performance computing node (3.80 GHz, 97.5 MB L3 cache, 96 cores). To evaluate the performance of S-LDSC, DESE, and scDRS in predicting the spatial localization of disease-associated cells, we utilized both a simulated spatial transcriptome dataset and multiple real-world datasets. For the real-world genetic association study (GWAS) component, we selected schizophrenia data from Vassily Trubetskoy et al.^37^ (n = 320,404). Additionally, we generated 6,000 synthetic GWAS samples for use with the simulated dataset. It is noteworthy that S-LDSC and scDRS are Python-based and lack multi-threading support, whereas DESE is implemented in Java and is multi-thread capable. To ensure a fair comparison, all methods were executed in single-thread mode.

Our simulated high-resolution dataset comprised a 100 × 100 spatial grid, representing 10,000 spots and encompassing 20,000 genes, analogous to data generated by technologies such as Stereo-seq.

We recorded runtime and peak memory usage for a high-resolution simulated dataset and four large-scale real-world datasets. The multi-threading capability of DESE was also evaluated by running it with 2, 4, and 8 threads.

## Code availability

The **SMECT** framework, analysis scripts, and curated resources are publicly available on GitHub at https://github.com/pmglab/smect.

## Data availability

The detailed information of the GWAS summary data is given in Supplementary Table 8. Mouse embryonic data: Stereo-seq, https://db.cngb.org/search/project/CNP0001543.

Mouse brain ST data: Stereo-seq, https://db.cngb.org/search/project/CNP0001543.

Macaque claustrum data: Stereo-seq, https://www.braindatacenter.cn/datacenter/web/#dataSet/details?id=1904344961189928962.

Human DLPFC ST data: https://research.libd.org/globus/.

## Extended data

**Extended Data Fig 1:**
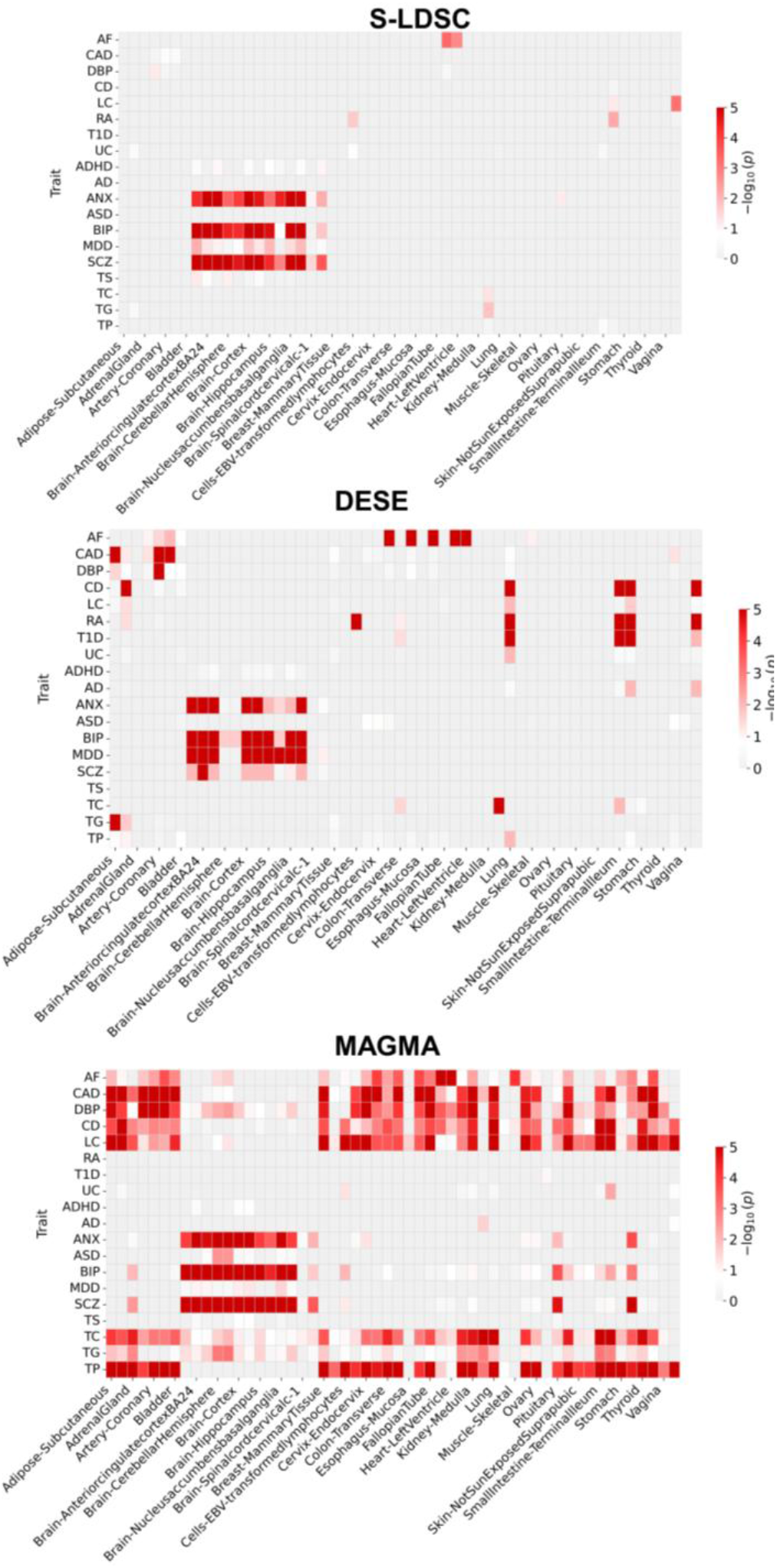
Positive control analysis validating genetic input quality using GTEx bulk RNA-seq data. Heatmaps display normalized enrichment scores (−*log*_10_ *p*) across 14 human traits (rows) and 49 tissues (columns) for three methods (S-LDSC, DESE, MAGMA), with intensity identically scaled (white to red: low to high significance).

**Extended Data Fig 2:**
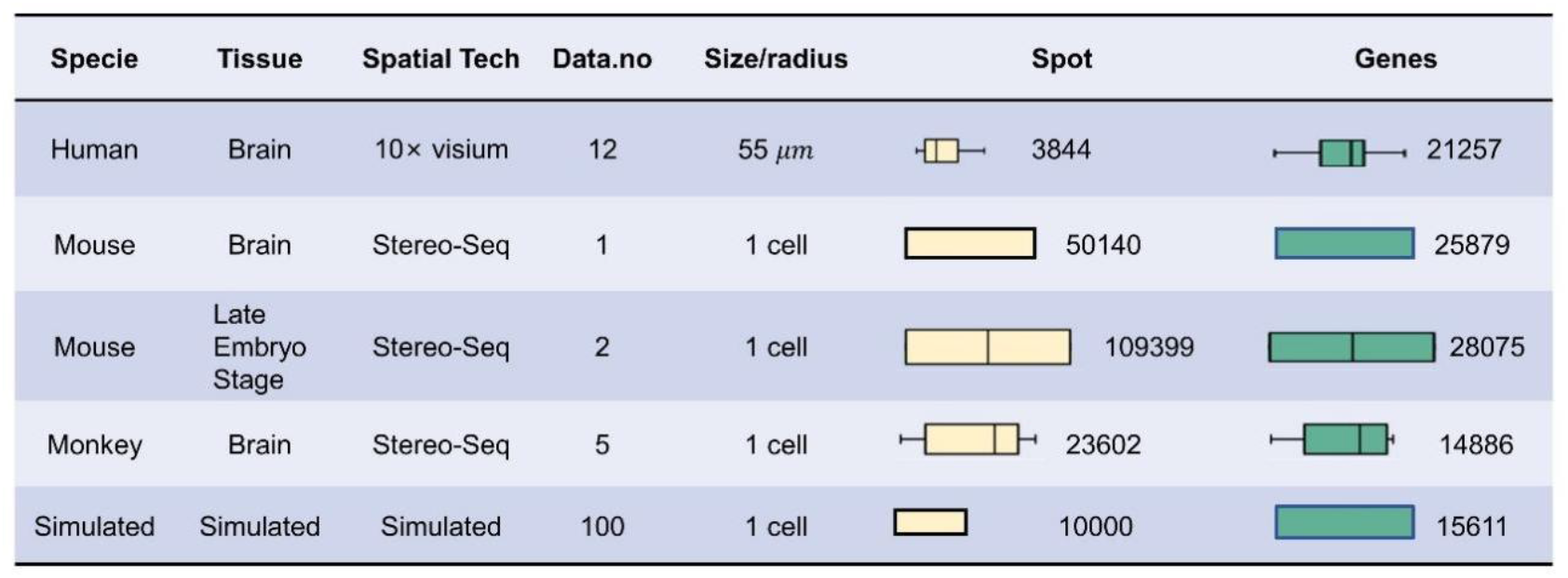
Characteristics of the real-world and simulated datasets used in this study. The table provides details including Species, Tissue type, Spatial Technology, Data number (Data.no), Size/radius of spots/cells, the number of Spots, and the number of Genes. Box plots illustrate the distribution of spots and genes for each dataset.

**Extended Data Fig 3:**
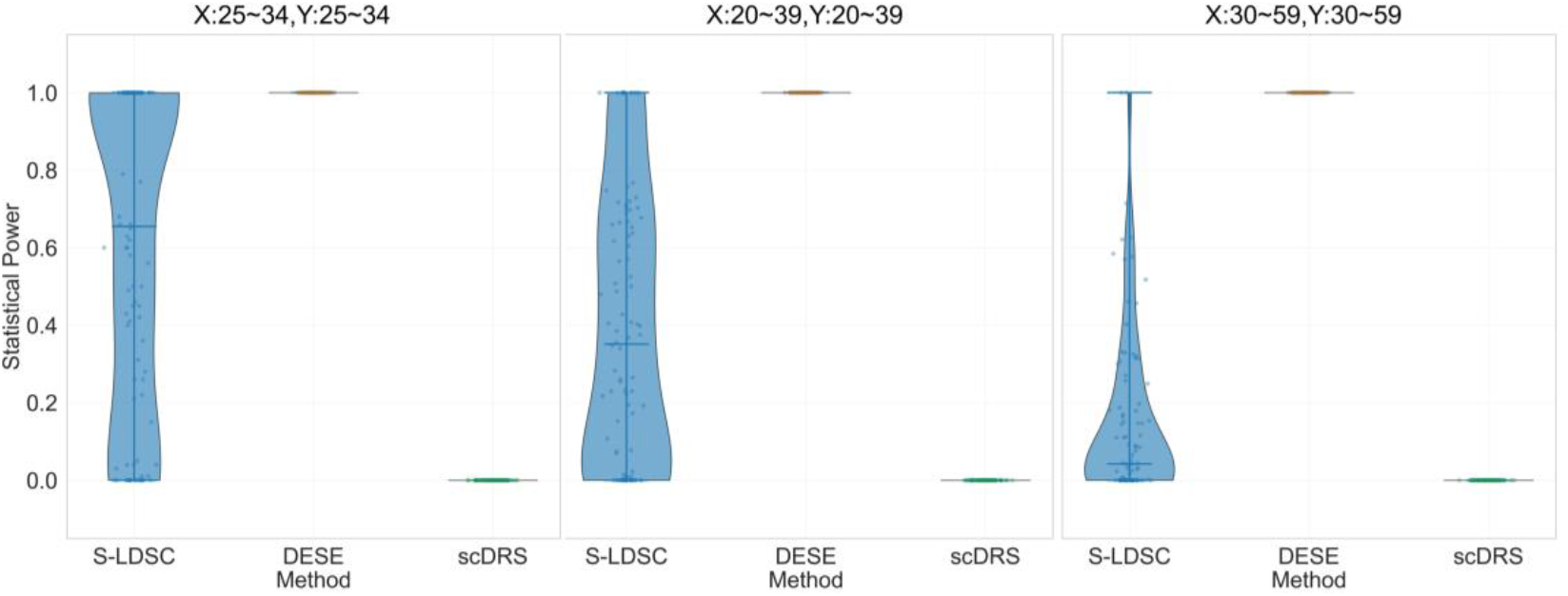
Violin plot of the statistical power of three methods under different simulation scenarios. These violin plots present the distribution of statistical power across three analytical methods (S-LDSC, DESE Method, scDRS) under three distinct simulation scenarios (differentiated by the simulated “true disease regions” in each subplot: Left subplot: Simulation scenario with X in 25∼34 and Y in 25∼34; Middle subplot: Simulation scenario with X in 20∼39 and Y in 20∼39; Right subplot: Simulation scenario with X in 30∼59 and Y in 30∼59). The y-axis in each subplot represents the “statistical power” of each simulated slice (ranging from 0 to 1), while the x-axis corresponds to the three methods.

**Extended Data Fig 4:**
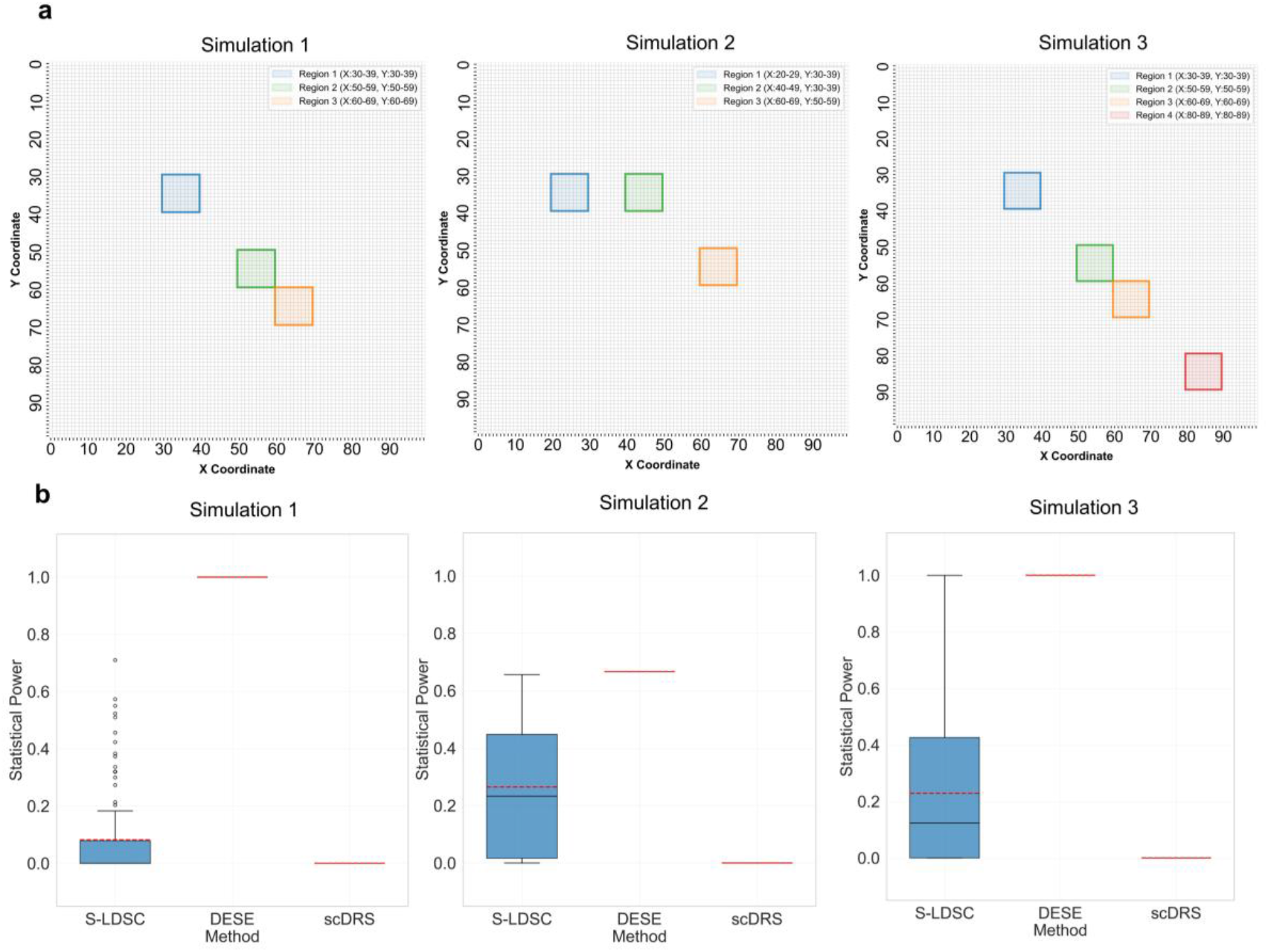
Statistical power results under different simulated scenarios in multiple regions. a. Panel a presents the spatial distribution of prespecified “true disease regions” across three simulation scenarios (Simulation 1, 2, and 3), where each subplot corresponds to one simulation scenario. Different colored squares represent the prespecified simulated “true disease regions”, and the legend clarifies the coordinate ranges for each region, intuitively demonstrating the spatial location differences of “true disease regions” among distinct simulation scenarios. **b**. Panel b shows the boxplots of statistical power distribution for a single slice under the three aforementioned simulation scenarios (each subplot corresponds to one simulation scenario). The boxplots depict the statistical power distribution characteristics of three methods (S-LDSC, DESE, scDRS) (the box represents the interquartile range), and the red horizontal line acts as a reference benchmark, enabling the comparison of statistical power performance of different methods across each simulation scenario.

