## Supplementary Methods for "SMECT: a framework for benchmarking post-GWAS methods for spatial mapping of cells associated with human complex traits"

#### Detailed Simulation Framework

##### Spatial Transcriptomics Data Simulation Model

We developed a computational model to synthesize large-scale, biologically realistic spatial transcriptomics datasets. The model generates gene expression counts on a two-dimensional grid of spots, providing explicit control over spatial expression patterns, sparsity, and count variability.

*Spatial Covariance Modeling.*

For genes with spatially localized expression, we modeled the spatial correlation using the Matérn covariance kernel. The correlation between two spots separated by Euclidean distance $d$ is given by:

$$K(d;\nu,\ell)=\frac{2^{1-\nu}}{\Gamma(\nu)}\left(\frac{\sqrt{2\nu}d}{\ell}\right)^{\nu}K_{\nu}\left(\frac{\sqrt{2\nu}d}{\ell}\right)$$

where $\nu$ is a smoothness parameter, $\ell$ is the characteristic length scale, $K_\nu$ is the modified Bessel function of the second kind, and $\Gamma$ is the gamma function. To ensure numerical stability, we approximated the kernel using its integral representation with Gauss-Laguerre quadrature ($n = 64$):

$$\kappa_{\nu}(r)\approx\frac{1}{2^{\nu-1}\Gamma(\nu)}\sum_{j=1}^{64}w_j\,x_j^{\nu-1}\exp\left(-\frac{\nu r^2}{4\ell^2 x_j}\right)$$

*Generation of Spatially Variable Gene Expression.*

The latent mean expression $\mu_{i,j}$ for gene $i$ at spot $j$ is modeled as a sum of a baseline expression level $\beta_i$ and contributions from $K$ expression hotspots:

$$\mu_{i,j}=\beta_i+\sum_{k=1}^{K}a_{i,k}\cdot K(\parallel \mathbf{x}_j-\mathbf{c}_k\parallel;\nu,\ell)$$

where $\mathbf{c}_k$ are the hotspot coordinates and $a_{i,k}$ are the amplitudes determining peak expression intensity. To model technical artifacts, the latent mean was set to zero with a probability $p_0$ (dropout), and the final integer count $Y_{i,j}$ was sampled from a Poisson distribution: $Y_{i,j}\sim \text{Poisson}(\mu_{i,j})$.

#### Generation of Simulated GWAS Summary Statistics

To create a ground truth for benchmarking, we designed a hierarchical model to simulate a quantitative phenotype whose genetic architecture is mediated by gene expression. This allows us to generate realistic GWAS summary statistics corresponding to our simulated spatial transcriptomics data.

*Stage 1: Simulating Gene Expression Levels.*

The expression level of gene $j$, $GE_j$, is the sum of its genetic value ($G_{GE,j}$) and an environmental component ($E_{GE,j}$). The genetic value is determined by its $k_j$ eQTLs:

$$G_{GE,j} = \sum_{i=1}^{k_j} X_{ji}\,\alpha_{ji}$$

The environmental component is drawn from a normal distribution, $E_{GE,j} \sim N(0, \sigma^2_{E_{GE,j}})$, with its variance determined by the gene expression heritability, $h^2_{GE}$:

$$\sigma^2_{E_{GE,j}} = \mathrm{Var}(G_{GE,j})\left(\frac{1 - h^2_{GE}}{h^2_{GE}}\right)$$

*Stage 2: Simulating the Final Quantitative Phenotype.*

The final phenotype $P$ is decomposed into three independent components: a pure genetic component ($G_{P,eQTL}$), a downstream non-genetic component ($C_{GE,env}$), and a residual component ($E_{p,resid}$):

$$P = G_{P,eQTL} + C_{GE,env} + E_{p,resid}$$

where $G_{P,eQTL} = \sum_{j=1}^{m} b_j\, G_{GE,j}$ and $C_{GE,env} = \sum_{j=1}^{m} b_j\, E_{GE,j}$. The heritability of the phenotype, $h^2_{P,eQTL}$, is defined as the proportion of variance explained by the pure genetic component, $\mathrm{Var}(G_{P,eQTL})/\mathrm{Var}(P)$. From this, we derive the variance of the final residual term:

$$\sigma^2_{E_{p,resid}} = \left[\mathrm{Var}(G_{P,eQTL})\left(\frac{1 - h^2_{P,eQTL}}{h^2_{P,eQTL}}\right)\right] - \mathrm{Var}(C_{GE,env})$$

*Constraint for a Valid Simulation.*

For the residual variance $\sigma^2_{E_{p,resid}}$ to be non-negative, the heritability of the final phenotype mediated by eQTLs cannot exceed the heritability of the intermediate gene expression traits. This imposes a critical constraint on the input parameters: $h^2_{P,eQTL} \leq h^2_{GE}$.

#### Simulation Design and Parameterization

We conducted simulations in two settings: (1) generation of a $100 \times 100$ spatial transcriptomic dataset, and (2) simulation of a large-scale population for a case-control study.

- **Gene Universe:** We simulated 20,000 genes, of which 500 were modeled with spatially variable expression patterns.
- **Disease Susceptibility Genes:** Among the 500 spatially variable genes, 250 were designated as true disease susceptibility genes.
- **Spatial Expression Parameters:** We set the smoothness $\nu = 1.5$, characteristic length scale $\ell = 5$, and kernel cutoff = $10^{-3}$. The baseline expression and dropout rate were 1.0 and 0.4, respectively. For power evaluation, a single expression hotspot for the 250 susceptibility genes was placed at grid location (30,30). For Type I error evaluation, no hotspots were included.
- **Genotype Data:** Genotypes for common variants (MAF > 0.05) were sourced from 490,541 subjects in the UK Biobank (UKB) cohort.
- **eQTLs and Expression Variance:** Each of the 250 susceptibility genes was influenced by 1 to 5 eQTLs, which collectively explained 25% of the variance in its target gene's expression ($h^2_{GE} = 0.25$).
- **Genetic Contribution to Liability:** The collective expression of all 250 susceptibility genes explained 25% of the total variance in disease liability ($h^2_{P,eQTL} = 0.25$). The weights $b_j$ were fixed to 1.
- **Disease Prevalence:** The population prevalence of the simulated disease was set to 2%.

*Case-Control Study and Association Analysis.*

From the simulated large-scale population, a case-control sample of 3,000 cases and 3,000 controls was drawn. Genetic association analysis was performed using a chi-square test to generate GWAS summary statistics.

### Detailed Statistical Analyses

#### Stability Analysis in Human Brain Tissue

For the robustness analysis of four replicate human DLPFC slices, we first derived spot-level p-values for schizophrenia association. We then aggregated these p-values to obtain a single p-value for each of the six cortical layers using the Cauchy combination test. The stability of predictions was quantified by calculating the pairwise Spearman's rank correlation coefficient of these layer-specific p-values across all four slices.

#### Spatial Autocorrelation Analysis

To quantify the spatial organization of disease-associated genetic signals, we performed Moran's I analysis on the adjusted p-values within the glutamatergic (Glu) neuron population from the adult mouse brain dataset. Moran's I was calculated using the formula:

$$I = \frac{n}{\sum_{i=1}^{n}\sum_{j=1}^{n} w_{ij}} \frac{\sum_{i=1}^{n}\sum_{j=1}^{n} w_{ij}\,(x_i - \bar{x})(x_j - \bar{x})}{\sum_{i=1}^{n}(x_i - \bar{x})^2}$$

where $n$ is the number of Glu-neurons, $x_i$ is the adjusted p-value for neuron $i$, $\bar{x}$ is the mean of the p-values, and $w_{ij}$ is the spatial weight between neurons $i$ and $j$. A spatial weights matrix was constructed using an inverse-distance scheme, with a cutoff distance applied to define local neighborhoods within a biologically relevant range. The matrix was row-standardized before calculation.

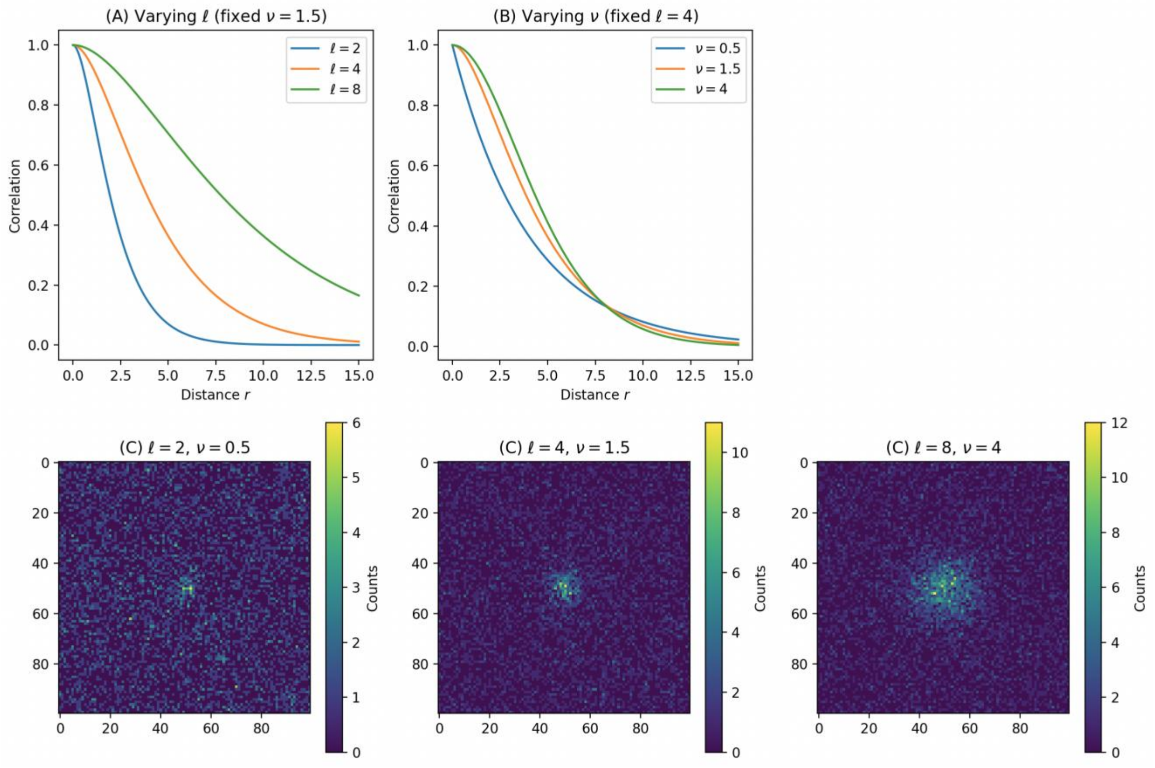


Fig. S1| Simulation of spatial transcriptomics data with tunable Matérn parameters. (A) Effect of varying the correlation length scale $\ell$ (with fixed $\nu$). Larger values yield broader spatial correlation. (B) Effect of varying the smoothness parameter $\nu$ (with fixed $\ell$). Larger values produce smoother spatial fields. (C) Simulated $100 \times 100$ spatial expression maps for different combinations, showing hotspot locations (bright central pixels) and the decay of expression intensity. Hotspots are generated by convolving their locations with a precomputed Matérn kernel, followed by zero inflation and Poisson sampling.

#### Statistical power under the alternative hypothesis

To assess the statistical power of each method, we utilized the 100 simulated datasets generated under the alternative hypothesis (i.e., containing a predefined disease region). For each simulation, we evaluated the method's ability to detect this region. Under a single disease hotspot, statistical power was quantified using a Gaussian-weighted scoring metric, defined as follows:

Let $(x_c, y_c)$ be the known central coordinate of the simulated disease region, and $\sigma = 1.1$ be the standard deviation for the Gaussian decay. The weight $w_d$ for a distance from the center is given by:

$$w(d) = e^{-\frac{d^2}{2\sigma^2}}$$

Where $d_i = \sqrt{(x_i - x_c)^2 + (y_i - y_c)^2}$ .For each simulation run, the power was assigned based on the following criteria:

$$Power = \begin{cases} 1.0 & if (x_c, y_c)\ exists\ and\ FDR < 0.05 \\ w(d_{min}) & if\ \exists (x_i, y_i) \in D_{disease}\ with\ FDR < 0.05 \\ 0 & otherwish \end{cases}$$

Under multiple disease hotspots, the power of a single slice is calculated as the proportion of significant spots within the disease area. The overall power is the average of all simulated slices.
